# Interannual surface water CO_2_ and O_2_ dynamics during fall in a small headwater lake

**DOI:** 10.1101/2025.06.30.659355

**Authors:** Brandon Blanchette, Morgan Botrel, Raoul-Marie Couture, Paul del Giorgio, Alice H. Parkes, Roxane Maranger

## Abstract

Lake metabolism is often quantified using continuous measures of dissolved oxygen (O_2_), where a 1: -1 stoichiometry with carbon dioxide (CO_2_) is assumed because of their roles in photosynthesis and respiration, respectively. However, many other physical, chemical, and biological processes decouple dissolved O_2_ and CO_2_ concentrations in lakes. Tracking departures from 1:-1 stoichiometry may provide insights into larger scale ecosystem functioning, particularly during fall when temperatures change and destratification occurs. Using continuous measures of both dissolved O_2_ and CO_2_ in a small temperate headwater lake, we looked at the interannual gas departure signals during fall over seven years. The beginning of fall, defined here as the start of leaf colour change, differed among years but coincided well with the onset of lake destratification and a shift in surface gas concentrations. Fall surface CO_2_ accumulation rates varied considerably, whereas O_2_ depletion rates were rather similar among years. Departure signals were broadly related to interannual differences in climate: more CO_2_ accumulated in the surface during the hottest-wettest fall compared to the coldest-driest one (0.81 and 0.37 µmol L^-1^ d^-1^, respectively), presumably from more catchment than hypolimnetic inputs. Lower CO_2_ accumulation occurred during years with prolonged hypolimnetic hypoxia potentially through enhanced CO_2_ consumption by methanogenesis. Other internal biological phenomena influenced fall departure signals, including a large metalimnetic oxygen peak, and higher fall surface primary production. We suggest gas departures during fall provide an integrative metabolic fingerprint for temperate stratified lakes, as well as insights into winter-priming conditions.

**Highlights:**

- Annual fall surface water CO_2_ accumulation rates vary more than O_2_ depletion rates.
- External fall inputs and internal processes influence gas departures differently.
- Fall gas departures may act as an integrative metabolic signal in temperate lakes.

## Introduction

Given its pivotal role in major biological processes and the increased availability of affordable, reliable *in situ* O_2_ sensors, continuous measures of dissolved oxygen (O_2_) are now commonly used to estimate whole lake ecosystem metabolism (Staehr and others 2010; Klanten and others 2023). Ecosystem metabolism involves the creation and use of energy through photosynthesis and aerobic respiration, where both processes respectively produce and consume O_2_ as well as consume and produce carbon dioxide (CO_2_). Although the two gases of these complementary co-occurring biological processes can be coupled in lake surface waters (Cole and Caraco 2001; Hanson and others 2006), they are not interchangeable in those assessments of ecosystem metabolism. Decoupling between O_2_ and CO_2_ has been observed over different time scales and across systems (Torgersen and Branco 2008; Stets and others 2009; Seybold and others 2021), and this decoupling in lakes can be a function of various biological, chemical, and physical processes (Vachon and others 2020; Vaziourakis and others 2025) such as chemolithotrophic microbial processes, chemical oxygen quenching, and wind driven mixing events, among others.

Direct and continuous measures of both O_2_ and CO_2_ gas concentrations in lake surface waters remain rare, and those that exist have focused on departure dynamics across systems during summer months (Laas and others 2016; Vachon and others 2020). These among-system studies provide useful information on the dominant ecosystem processes that influence the gases (Crawford and others 2014; Vachon and others 2020). Yet interannual assessments of paired O_2_-CO_2_ measures may be particularly insightful in providing a more integrative metabolic signal or “fingerprint” within a single system (Bernhardt and others 2018), especially during dynamic shoulder seasons such as fall, where changing environmental conditions can influence surface gas concentrations differentially.

Fall, in boreal and temperate regions, is a season characterized by widespread climatic and ecosystemic changes in lakes as well as their surrounding catchments (Kalff 2002; Woolway 2023). During fall, particularly in temperate regions, the leaves of deciduous trees begin to change colour and fall to the ground in response to decreases in air temperature, marking the end of evapotranspiration (Lim and others 2007; Xie and others 2018). As the hydrology of lakes in these regions are highly regulated by their forested catchments (Allan and Roulet 1994; Buttle and others 2004), the end of evapotranspiration results in an increased hydrologic exchange and a higher run-off ratio compared to summer (Kalff 2002). This results in CO_2_ rich waters entering the lakes from their catchment (Humborg and others 2010; Weyhenmeyer and others 2015), particularly in years when precipitation is high (Raymond and Saiers 2010; Einola and others 2011; de Wit and others 2018). Fall also marks the moment of lake destratification in response to decreasing air temperature and increasing winds, where potentially anaerobic hypolimnetic bottom waters rich in CO_2_ and poor in O_2_ mix with those of the surface (Kalff 2002). The overall volume of anaerobic bottom waters however can differ among years, as climatic conditions during the open water season influence hypoxic duration (Couture and others 2015; Bartosiewicz and others 2019; LaBrie and others 2023), and hypolimnetic extent (Woolway and others 2021; Carey 2023). Furthermore, interannual changes in the relative inputs during fall expressed by the O_2_-CO_2_ departure signal may result in variable winter conditions that will influence community structure (Hampton and others 2024). By measuring how the changes in daily estimates of O_2_ and CO_2_ track each other over time (Bernhardt and others 2018), we can examine how the metabolic fingerprint varies as a function of different climate drivers, which can vary quite significantly across years.

In this study, we explore the interannual variability of continuously measured paired O_2_-CO_2_ in a small headwater lake during the fall for seven years. We hypothesize that CO_2_ rich and O_2_ poor inputs from the watershed due to reduced water retention capacity combined with hypolimnetic inputs during destratification will lead to a strong decoupling between these gases in the surface waters during fall. We expect both these internal and external controls to lead to patterns of increased CO_2_ and decreased O_2_ during this period, and anticipate that the response will differ among years as a function of both the weather and relative inputs of the gases. Departure differences could reflect broader integrative lake dynamics influenced by interannual variability of climate, and within lake processing, but to our knowledge this characterization has yet to be done. We anticipate an increased CO_2_ accumulation and O_2_ depletion in lake surface waters in years with higher precipitation during fall and in years where the lake experienced a greater hypolimnetic extent or hypoxic duration. Interannual differences in the relative contributions of both gases from these external and internal controls will allow us to explore differences in the annual metabolic fingerprint.

## Methods

### Study site and data collection

The study was carried out in Lac Croche (45°59’32” N 74°00’18” W), a small (0.179km^2^) oligotrophic headwater lake with a pristine forested catchment dominated by sugar maple (*Acer saccharum)* and yellow birch (*Betula alleghaniensis*) located at the Université de Montréal field station on the Canadian Shield, in Québec, Canada. Lac Croche has a small watershed (1.07 km^2^), and its waters are primarily renewed from groundwater inputs and surface runoff. The lake has three connected basins that are circumneutral in terms of their pH (6.6 ± 0.4), it stratifies seasonally, and is ice-covered during winter. Using sensors on a floating platform deployed at the deepest point of the lake (maximum depth ∼11m) in the eastern downstream basin (Figure 1S), surface O_2_ and CO_2_ gas dynamics were measured during the open water season from approximatel yearly May to early November in 2015 to 2022, except in 2020 when there was no deployment due to the COVID-19 pandemic. Our study focused on understanding the gas departures during fall, defined as the moment when leaves of the deciduous trees in this temperate mixed forest started to change colour, reflecting a broader ecological moment as opposed to a set calendar date.

### Continuous lake measurements

The autonomous floating platform was equipped with a variety of underwater sensors, the focus being on temperature, CO_2_, and O_2_ in this study. Surface water pCO_2_ was measured at a depth of 0.5m using a submerged infrared CO_2_ sensor (Vaisala GMP343, Finland, accuracy ± 0.5%). This sensor was covered with a diffusion membrane (PTFE) that allows the gas to equilibrate without having to pump air. Dissolved oxygen optical sensors (T-Chain RS 232/485, Precision Measurement Engineering Inc., USA, precision of ±0.3 mg L^-1^ or ±5% of the measurement, whichever is larger) were deployed at 0.5m, 5.5m and 8.5m depths, while temperature (T-Chain RS 232/485, Precision Measurement Engineering Inc., USA, precision of +0.010 °C) was recorded at about every meter between 0.5m and 10.5m with slight variations in the precise depth among years. In 2015, a copper screen was placed on the CO_2_ probe to prevent biofouling that led to a decrease in the absolute concentrations due to the formation of copper carbonate (CuCO_3_). Relative changes in CO_2_ values however were unaffected, and the copper screen was not used in subsequent years.

### Gas measurements from sources

To measure the contribution of gases from catchment sources relative to those from the hypolimnion, water samples were collected from piezometers placed in the riparian and upland zones in early November 2023, and from the hypolimnion weekly through the fall in 2022. Water from the piezometers were collected by gently filling 100 ml into three 140 ml syringes. The headspace was created by introducing 40 ml of Ultra Zero air (containing no CO_2_) from a foil bag (Supel-Inert, Supelco). The syringes were vigorously shaken for two minutes to equilibrate the dissolved gases with the headspace, then the headspace was injected into 9 ml pre-evacuated glass vials. Water from the hypolimnion was collected at 9 m using a Van Dorn water sampler, and gently transferred to a 1.1 L glass bottle where gas samples were collected using the headspace equilibration technique on site. Once hermetically sealed, 0.12 L of water from the 1.1 L bottle was replaced with ambient air by pulling water using a syringe. The bottles were vigorously shaken for two minutes, and three replicate headspace samples were collected and injected into 9 ml pre-evacuated and sealed glass vials. One ambient air sample was taken at each site to measure the partial pressures of gases in the atmosphere and in the pre-equilibration headspace for the bottle samples.

Headspace samples were analyzed for CO_2_ by gas chromatography (Shimadzu GC-2014— Shimadzu Scientific Instruments, Columbia, United States) equipped with a TCD. Three different concentrations (200 ppm, 1503 ppm, 10 000 ppm) were used for our calibration curves (precision ±5%). Oxygen concentrations in the lake were measured using a multiparameter probe (YSI 650 MDS - Yellow Spring Instruments, Ohio, United States) and water samples from the riparian and upland soils were collected in 60 ml biological oxygen demand (BOD) bottles using a peristaltic pump. One ml of manganese chloride and alkaline iodide were subsequently added to the samples and thoroughly mixed and sealed. Oxygen content was then determined using the Winkler titration method (Winkler 1888).

### Climate and fall timing

Climate information extracted from the European Centre for Medium-Range Weather Forecasts (ECMWF), ECMWF Reanalysis v5 (ERA5) climatic reanalysis dataset (Hersbach and others 2017) was used for air temperature, barometric pressure, humidity, precipitation, wind speed, cloud cover and global radiation. This dataset uses forecast models and data assimilation systems to reanalyze archived observations creating a global 31 km cell grid with hourly climatic monitoring data. For this study, we used daily means of climate data from May 14^th^, 2015, to December 31^st^, 2022. The ERA5 climate data was required because data from the closest weather station (Government of Canada 2023) to Lac Croche and *in situ* sensors were incomplete. We, however, validated the ERA5 dataset against data from local weather stations for air temperature, wind and precipitation where possible (Figure 2S).

The start of fall, defined as the moment of leaf colour change, was determined using images from a PhenoCam Network camera as part of a broader network of forest sites across the US and Canada (Milliman and others 2019). The camera is located in the forest on the south shore of the Lac Croche study basin and takes hourly images during daytime (∼14 images per day). For each year, images were inspected from early September, and the beginning of fall was noted as the day where visible deciduous tree leaves changed colour. This identification was straightforward and done by a single person minimizing potential uncertainty in identifying the moment colour change began. End of the study period was the date of buoy removal which occurred between the 301^st^ and 309^th^ Julian days.

### Modeling lake temperature dynamics

To fill gaps in our water temperature time-series, we simulated water temperature using the 1D process-based lake model MyLake v 1.2 (**M**ulti-**y**ear **L**ake simulation model; Saloranta and Andersen 2007). MyLake is a MATLAB code that simulates water column stability, daily distribution of heat, and other variables using a stacked layered geometry consisting of mixed horizontal layers (Saloranta and Andersen 2007). Process-based models provide advantages over empirical ones as they explicitly consider in-lake phenomena, such as wind-induced mixing, heat capacity, and solar radiation penetration rather than relying only on daily stochastic weather progression (Fang and Stefan 2009). MyLake uses lake bathymetry and assumes lateral homogeneity of water temperature. MyLake has been successfully used to study lake hydrodynamics and biogeochemistry including the impact of climate on lake thermal regimes (Read and others 2011; Dibike and others 2012; La Fuente and others 2022), dissolved oxygen dynamics (Couture and others 2015) and ice formation (Gebre and others 2014). When used solely to simulate water temperature time-series, MyLake – as is the case for other 1D process-based lake models – performs very well with minimal calibration using its default parametrization, bathymetry and observed weather as forcings (Côté and others 2023). For this application, only the eastern basin of Lac Croche, where the sensors were located was modeled. The model was forced by a time series of daily meteorological input data provided by the ERA5 dataset.

### MyLake model parameter estimation

Parameter values were estimated using manual calibration, by an iterative process whereby the values of each parameter were changed independently while maintaining all others constant and evaluating the corresponding goodness of fit. The parametrization was performed against temperature data from the years 2016-2019, i.e. the time series where all sensors were at constant depths (Figure 3S). Specifically, the parametrization was done for 0.5m, 1m, 4.5m, 6.5m, 8.5m and 10.5m depth. Afterwards, the model was validated with the time-series for 2015, 2021, and 2022 (Figure 3S). The temperature profiles in 2015 were not measured at the same depths as in other years, while 2021 and 2022 ensured appropriate model performance (Figure 4S). Goodness-of-fit is reported as R^2^, to quantify the linear relationship between the modeled and measured data, and root mean square error (RMSE), to determine an overall average error between each data pair.

### Gas and environmental variable calculations

To compare relative changes in gas concentrations through the fall and among the different years, the CO_2_ and O_2_ gas concentration deviations from saturation were calculated and expressed as Δgas in µmol L^-1^ where Δgas = Conc_water_ – Conc_saturation_. Gas concentrations in the water were corrected for temperature and pressure to accurately represent the deviations from saturation as it changed over the season. To determine ΔCO_2_, the global average air concentration for each year was used as the saturation level (Lan and Thoning 2023). O_2_ saturation and ΔO_2_ were determined using the LakeMetabolizer package (Winslow and others 2016). The metrics for the O_2_: CO_2_ departure space of ecosystem quotient, offset, and stretch were subsequently calculated based on the cloud of points formed by each year (Figure 1; Vachon and others 2020). The ecosystemic quotient (EQ) is calculated as where the slope is a type II linear regression of the ΔO_2_: ΔCO_2_ cloud. EQ reflects the integrated stoichiometry of the ecosystem. The offset of the cloud is calculated as the sum of the CO_2_ and O_2_ departure averages and characterizes the extent or relative role of different processes such as external inputs and anaerobic respiration. The stretch is the major axis length of the 95% covariance error ellipse and characterizes the variability around the dominant process.

**Figure 1.**
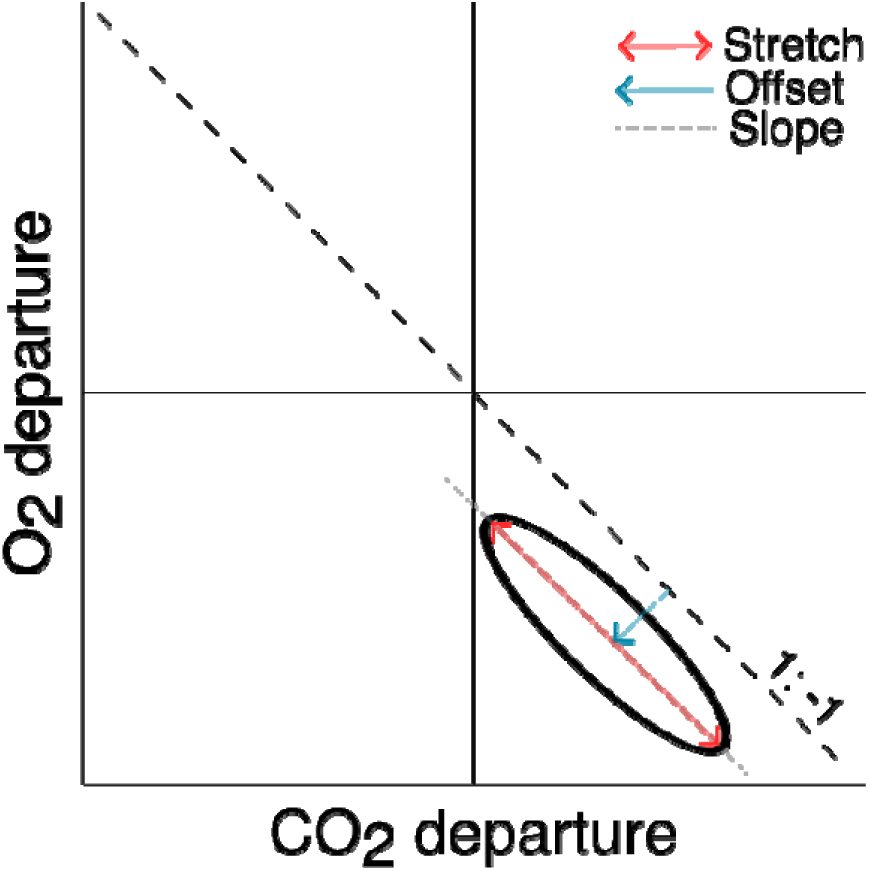
Conceptual figure of metrics to characterize the O_2_ -CO_2_ departure space. The black ellipse represents an example cloud of points (adapted from Vachon and others 2020).

Continuous water temperature measurements from the floating platform and bathymetric data were used to calculate the lake thermal stability as the Schmidt stability (St, J m^-2^) using the Lake Analyzer Package (Read and others 2011; Winslow and others 2019). This metric represents the areal amount of energy required to completely mix the lake. To estimate the period when lake bottom waters had low oxygen concentrations, hypoxic duration was calculated as the cumulative number of days since dissolved oxygen concentrations went below 2.0mg/L (LaBrie and others 2023). To determine the contribution of water from the hypolimnion during fall destratification, hypolimnetic volume was calculated using 1-m bathymetric curves.

Finally, in order to estimate the total mass amount of CO_2_ reaching the surface from deeper layers during fall destratification, we used a simple mixing model where we calculated the CO_2_ concentration in the adjacent layer mixing with the surface by using the change in CO_2_ surface concentrations and total volume pre and post mixing. We then estimated the total amount of CO_2_ as a function of the volume of that layer and summed the total CO_2_ content from those adjacent layers at each time step of mixing over the destratification period. Layer and mixing volumes were determined with MyLake temperature profile estimates and changing surface ΔCO_2_ from the buoy. This exercise was done for the two contrasting years of 2015 and 2017, where more daily CO_2_ measurements from the surface were available from the buoy for the entire fall season, allowing for more accurate estimates.

### Statistical analyses

To assess the daily CO_2_ accumulation and O_2_ depletion rates of the fall season across years, we performed generalized least square (GLS) regressions (Pinheiro and others 2023). This approach accounts for the temporal autocorrelation structure of the data. We compared two models: one incorporating an autoregressive structure of order one (AR-1) and another one without (Table 1S and Table 2S). Best models were selected to minimize Akaike information criterion (AIC) and the GLS with the auto-regressive model was always a better fit, except marginally less so in 2019. Differences in these slopes were assessed using ANCOVA and post-hoc Tukey significant difference test (TukeyHSD; Table 3S and Table 4S). The O_2_: CO_2_ departure slopes were calculated using a type II linear regression for each year. A principal component analysis (PCA) was performed to describe the correlation between CO_2_ accumulation rate, O_2_ depletion rate, EQ and environmental variables (Figure 6; Le and others 2008). All statistical analyses and graphs were performed in R 4.3.0 (R 4.3.0 - R Core Team, 2023). AI was used to help with writing code to generate figures.

## Results

The start of fall, which we defined as leaf colour change of deciduous trees in this temperate region, was variable among years, with the median start day of leaf colour change being on September 14^th^ but differing by up to 26 days, beginning as early as September 1^st^ in 2022, and as late as September 27^th^ in 2018 (Table 5S). The modelled temperature profiles showed clear thermal stratification patterns throughout the summers (Figure 2) with lake destratification rather consistently coinciding with the start of leaf colour change, albeit to a lesser extent in 2017 and 2022. In 2017, a slight restratification in early fall occurred (Figure 5S) when air temperatures were unusually high at >20 °C for five consecutive days, and as a result average air temperature in 2017 was higher in fall compared to other years (Table 5S). For 2022, fall began much earlier than usual (on September 1^st^ versus the median of September 14^th^), however despite the earlier onset, thermal lake stability began to decrease precisely at that moment like in other years (Figure 5S). As such, the start of fall appears to coincide with the onset of destratification, and a change in lake thermal dynamics. Full mixing of the water column was captured by the buoy in 2015 and 2018 on October 22^nd^ and October 18^th^ respectively, but in other years the buoy was removed for winter before turnover was complete. The turnover in these two years was comparatively early as the buoy was always removed around the last week of October or first week of November.

**Figure 2.**
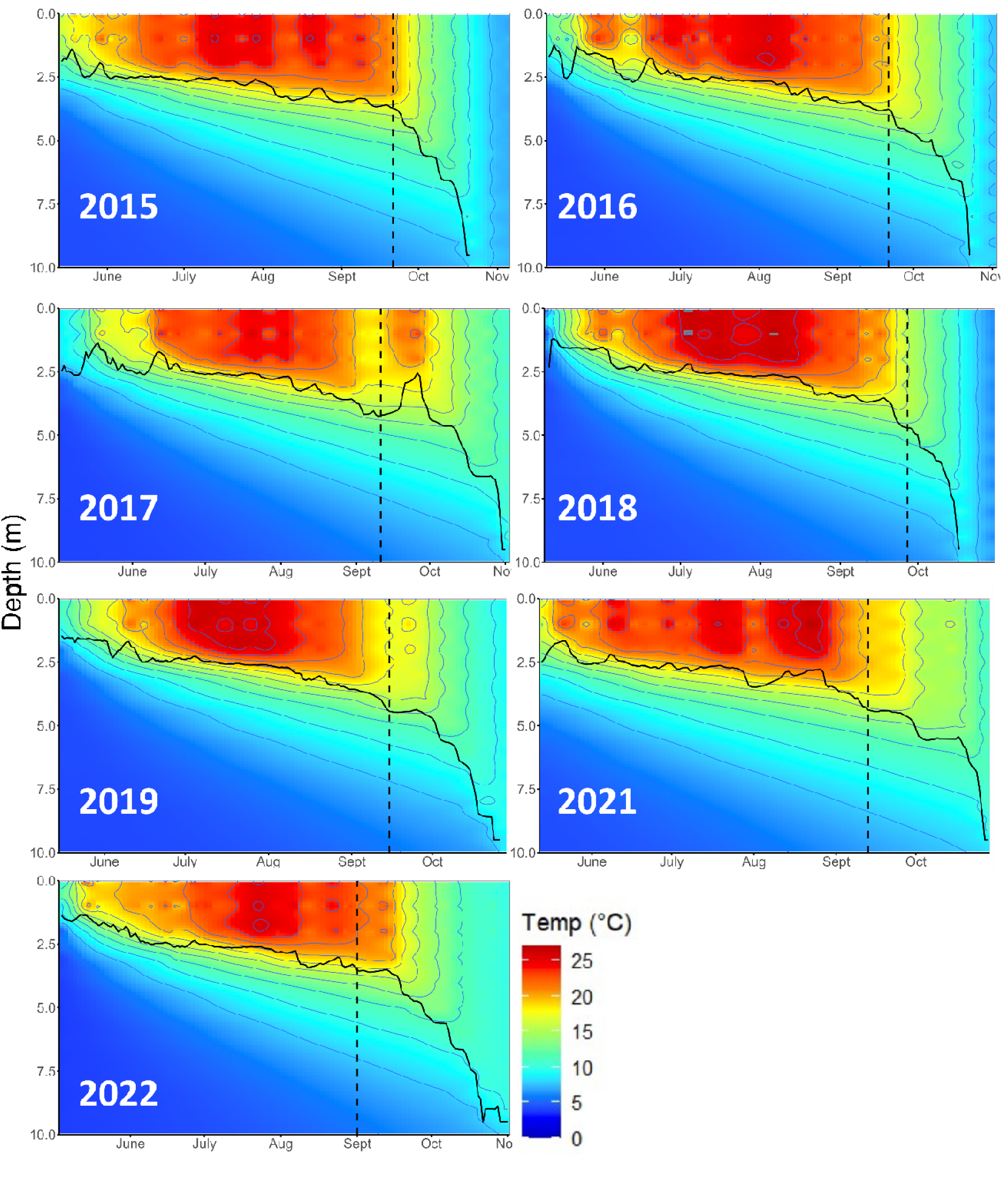
Modelled annual lake thermal stratification. Start of fall, defined as the start of leaf colour change, is represented by the black dashed vertical lines and thermocline depth is represented by the black continuous line. Dots represent dates with buoy data.

### Interannual variation of environmental variables

Environmental conditions during fall were characterized by a strong positive correlation between the cumulative rain volume and mean daily air temperatures across years (*r* = 0.96, Figure 3a). We observed an ∼8 °C difference in average temperatures and a twofold range of cumulative rain ranging from 13 to 31% of lake volume between extremes. Fall conditions in 2015 and 2018 were cold and dry, with mean air temperatures of 7 °C and 5 °C while receiving the lowest proportion of cumulative rain at 19 % and 12 % of lake volume (Table 5S), respectively. In contrast, 2017 was the hottest and wettest year with a mean fall temperature of 13 °C and cumulative rainfall of 31% of lake volume. Environmental conditions in 2019, 2021, and 2022 were more similar to 2017 while 2016 was more intermediate between the hot-wet versus cold-dry years. Because we defined the fall period as the start of leaf colour change until buoy removal, the duration of fall differed among years (mean 44 days, range 31-56 days). To verify if the interannual variability in environmental conditions was due to the different start dates and lengths in the fall period, these were recalculated using an average start date for a fixed fall duration among years. Regardless of how we define the fall period, the interannual differences in environmental conditions were similar. Therefore, we opted to use the start of leaf colour change to define fall environmental conditions. Cold years (2015, 2018) experienced higher average wind speeds and generally had the lowest cumulative radiation as compared to hotter years (2017, 2021; Figure 3b and d). Thermal stability loss, however, did not show a trend with cold to warm fall seasons, possibly because it is an integration of both wind speed and cumulative radiation (Figure 3b). Hypoxic duration throughout the ice-free period did not show a trend with cold to warm fall seasons (Figure 3e), possibly because it is determined by both the onset of thermal stratification in spring as well as the timing of destratification in fall.

**Figure 3.**
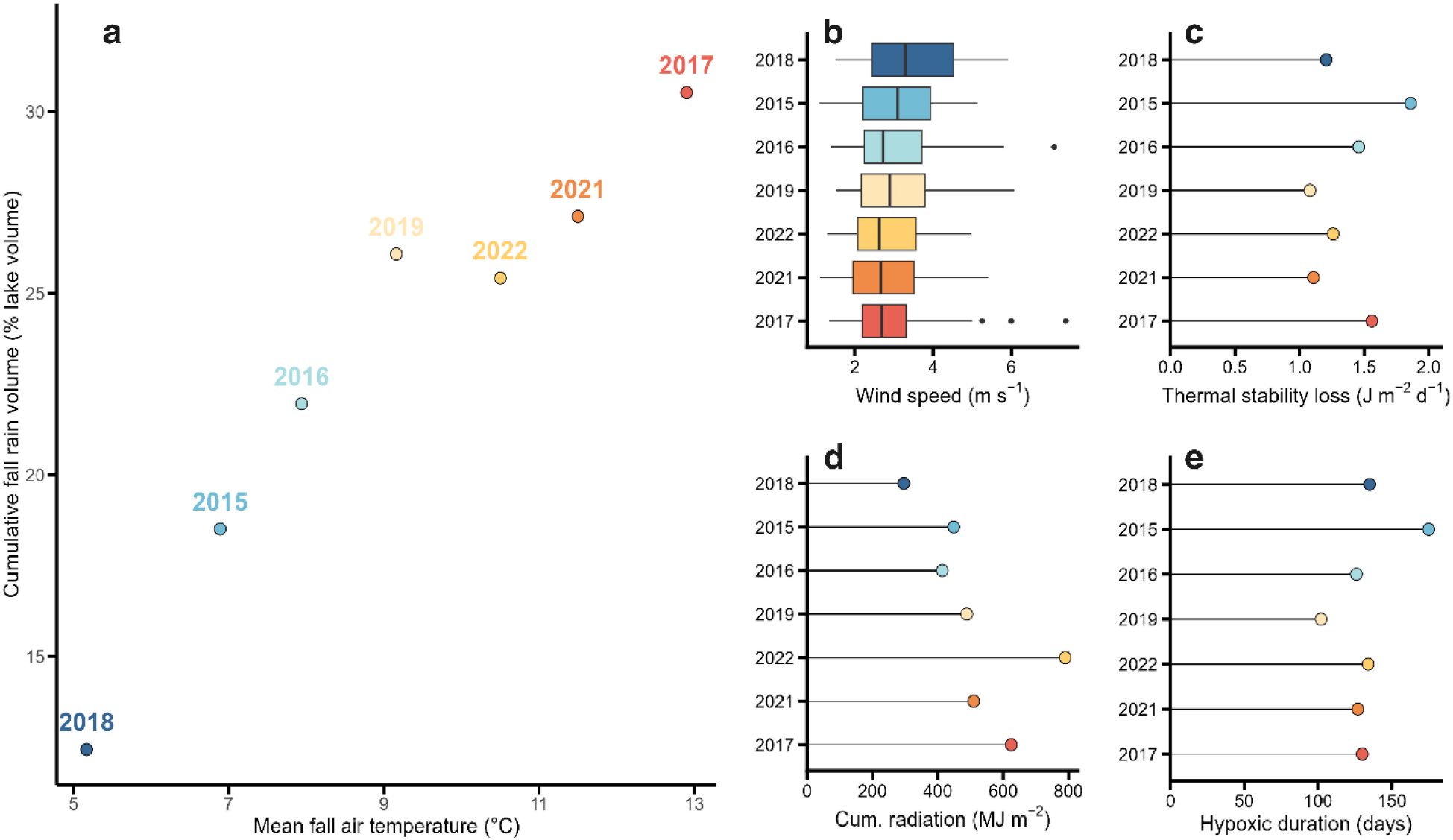
Interannual variation of environmental variables during fall (a-d) and summer (e) of 2015-2019, 2021-2022. Dot and boxplots are ordered by increasing air temperature using a cold to warm color gradient. Hypoxic duration is calculated over the course of the whole summer until mixing occurs.

### Carbon dioxide and dissolved oxygen dynamics

Surface water ΔCO_2_ and ΔO_2_ patterns observed during the summer (June 1^st^ – start of fall) were similar across years (Figure 4a). ΔCO_2_ was relatively stable over the course of the summer whereas ΔO_2_ dynamics were comparatively more variable within any of the years. ΔO_2_ was mostly stable between June and mid-August and slowly started to decrease gradually from that point forward. When looking at fall specifically, the surface concentrations of both gases changed dramatically and displayed either an abrupt increase (ΔCO_2_) or decrease (ΔO_2_, Figure 4a). There was significant interannual variability in CO_2_ accumulation rates among years (ANCOVA, *p* < 0.001), measured as the slope of the daily change in ΔCO_2_ concentration during fall, although some pairs of years behaved similarly and corresponded generally well to the cold-dry versus hot-wet distinction in environmental conditions across years (Figure 4b, Table 1). The cold-dry fall seasons (2018 and 2015) had the lowest CO_2_ accumulation rates of 0.37 and 0.55 µmol L^-1^ d^-1^ respectively, followed by rates that were higher and similar across 2021, 2022, and 2019, with the hot-wet fall season of 2017 being comparatively higher (Table 1). Finally, the rate of CO_2_ accumulation of 1.40 µmol L^-1^ d^-1^ in the intermediate temperature year (2016) was significantly higher than all other years (Table 1, Figure 4b and Table 3S). In comparison to CO_2_, the O_2_ depletion rates were more similar across years with only 2022 being significantly lower than some other years (0.62 µmol L^-1^ d^-1^, Table 1, Figure 4c and Table 4S).

**Figure 4.**
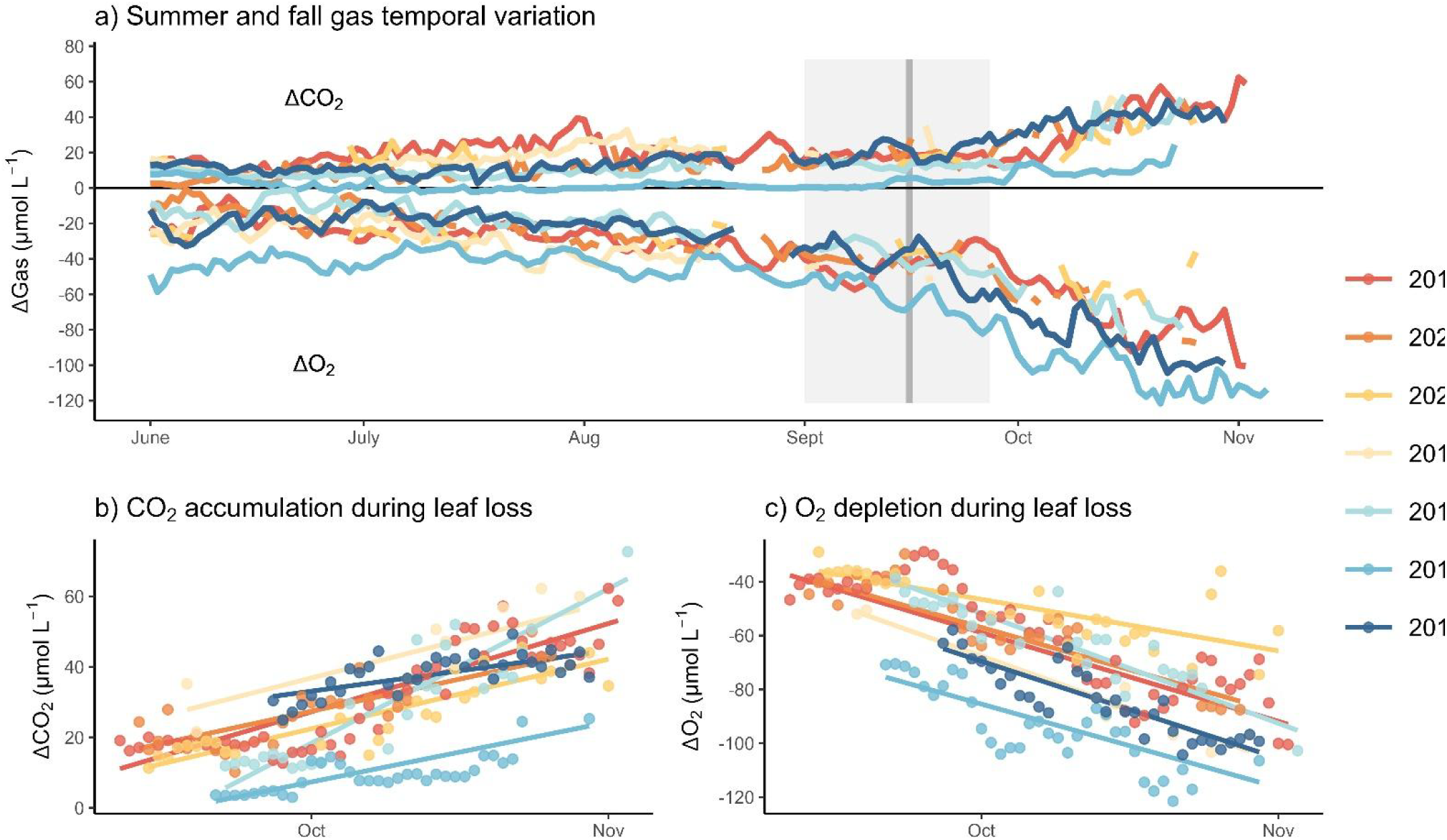
Time series of average daily temperature-corrected surface water ΔCO_2_ and ΔO_2_ patterns across years (a), CO_2_ accumulation (b) and O_2_ depletion (c) as a function of time during fall. The grey line in Figure a represents the mean start of leaf colour change while the grey box defines the range. Hot (red) to cold (dark blue) color gradient represents decreasing average air temperatures during fall periods.

**Table 1.** Interannual slope coefficients (μmol L^-1^ d^-1^) and variance explained (R^2^) by general least square regression models with autoregressive structure of order one for surface water ΔCO_2_ accumulation and ΔO_2_ depletion over time during fall for each year. Years ordered from coldest and driest to hottest and wettest. *Sample size (n) of 34 for ΔCO_2_ and 46 for ΔO_2_ in 2015. Diff. indicates pairwise significant difference at p < 0.03 determined from a post-hoc Tukey test following an ANCOVA.

| Environmental conditions | Year | $\Delta\text{CO}_2$ | | | | $\Delta\text{O}_2$ | | | | n |
| --- | --- | --- | --- | --- | --- | --- | --- | --- | --- | --- |
| | | Slope | $R^2$ | Diff. | range ($\mu\text{mol L}^{-1}$) | Slope | $R^2$ | Diff. | range ($\mu\text{mol L}^{-1}$) | |
| Cold & dry | 2018 | 0.37 | 0.58 | A | 31 – 44 | -1.16 | 0.78 | A | -104 – -65 | 34 |
| Cold & dry | 2015 | 0.55 | 0.51 | A | 3 – 18 | -0.99 | 0.75 | A | -122 – -77 | 34*, 46 |
| Intermediate | 2016 | 1.4 | 0.81 | C | 7 – 62 | -1.32 | 0.81 | A | -91 – -43 | 26 |
| Hot & wet | 2019 | 0.69 | 0.79 | AB | 26 – 60 | -1.26 | 0.94 | A | -103 – -51 | 9 |
| Hot & wet | 2022 | 0.64 | 0.77 | AB | 13 – 42 | -0.62 | 0.56 | B | -67 – -36 | 29 |
| Hot & wet | 2021 | 0.59 | 0.71 | AB | 16 – 44 | -1.02 | 0.83 | A | -86 – -35 | 26 |
| Hot & wet | 2017 | 0.81 | 0.78 | B | 9 – 53 | -1.06 | 0.77 | A | -90 – -30 | 53 |

The interannual variability of the CO_2_ accumulation and O_2_ depletion rates during fall were reflected in the CO_2_: O_2_ departure space (Figure 5), and the slopes for this relationship varied considerably among years ranging from -0.9 to -3.3 (Table 2). Metrics related to this departure space provide insights on the integrative metabolic response at the ecosystem level, where moles of O_2_ lost based on the ranges where up to 3 times more than CO_2_ gained (Table 2). Variability in the annual ecosystem quotient (EQ), which is the number of moles of CO_2_ produced per moles of O_2_ lost, shows how the integrative ecosystem stoichiometric response differs across years. The cold-dry years produced three times fewer CO_2_ relative to the amount of O_2_ lost (based on ranges), whereas several of the warm-wet years had similar EQs (Table 2). This suggests that beyond temperature and precipitation, other environmental drivers were acting on the integrated gas signal expressed during fall. The offset, or the sum of the average CO_2_ and O_2_ departures was similar for most years, ranging from -24.2 µmol L^-1^ to -35.9 µmol L^-1^, but the offset was greater for the cold-wet years 2015 and 2018 at -84.7 and -46.4 µmol L^-1^, respectively, where surface water CO_2_ concentration ranges were considerably lower (Table 2). The difference in the offset between these two years is likely due to the copper meshing, but the relative change remains robust. The stretch is the major axis length of the 95% covariance error ellipse, and along with the offset, captures the shape and position of the data clouds. This metric generally describes the strength of the metabolic signal driving these gas departures. Clustering of the stretch shares similarities to other data presented (Table 2) with lowest stretch values in cold-dry years, suggesting that gas co-evolution is less dynamic in those years, whereas hot-wet years (2017 and 2019) had higher stretch values, suggesting more temporal variability in the gas dynamics.

**Figure 5.**
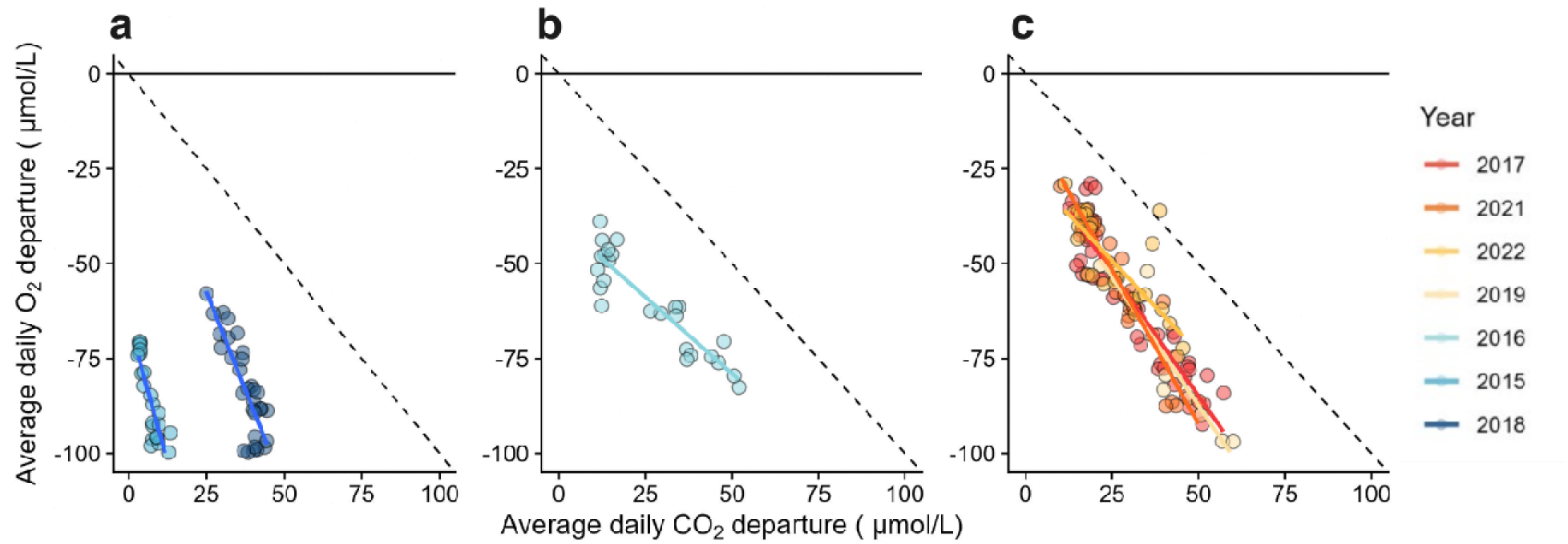
CO_2_ and O_2_ departures from atmospheric equilibrium during fall grouped by a) cold years, b) intermediate year, c) hot years. The dotted line represents the 1:-1 ratio of O_2_:CO_2_ departures.

**Table 2.** O_2_/CO_2_ slope and departure metrics, where EQ represents ecosystem quotient (1/slope) and values for offset and stretch are reported in µmol L^-1^. Years ordered from coldest and driest to hottest and wettest.

| Environmental conditions | Year | Slope<br>($\mu\text{M O}_2/\mu\text{M CO}_2$) | EQ<br>($\mu\text{M CO}_2/\mu\text{M O}_2$) | Offset<br>($\mu\text{mol L}^{-1}$) | Stretch<br>($\mu\text{mol L}^{-1}$) |
| --- | --- | --- | --- | --- | --- |
| Cold & dry | 2018 | -2.6 | 0.38 | -46.4 | 69.6 |
| Cold & dry | 2015 | -3.3 | 0.3 | -84.7 | 75.2 |
| Intermediate | 2016 | -0.9 | 1.12 | -33.3 | 108.8 |
| Hot & wet | 2019 | -1.5 | 0.68 | -35.9 | 111.2 |
| Hot & wet | 2022 | -1.3 | 0.77 | -24.2 | 76.1 |
| Hot & wet | 2021 | -1.8 | 0.57 | -28 | 95.5 |
| Hot & wet | 2017 | -1.4 | 0.71 | -29.2 | 120.7 |

### Potential drivers of interannual variability of CO_2_, O_2_ rates and ecosystem quotient

We used a PCA to better elucidate which drivers were associated with the observed interannual variability in EQ, CO_2_ accumulation and O_2_ depletion rates beyond temperature and precipitation (Figure 6; Table 5S). The first two principal components explained a large portion of the variability (73.3%), with the first component related largely to external environmental drivers, accounting for 42% and the second more related to internal lake dynamics, explaining 31.3%. The EQ was located between both axes, suggesting that it was an integrative metric influenced by both external and internal drivers subject to interannual variation. The cold-dry falls of 2015 and 2018 had the lowest overall EQs and CO_2_ accumulation rates, but these were explained by different internal and external drivers. Both years started with low O_2_ concentrations in their bottom waters, however 2015 had less than half the concentration of 2018 at 0.78 versus 1.74 mg L^-1^ respectively (Figure 6S). This combined with 2015 being strongly stratified with the hottest summer on average (Table 5S) resulted in that year having the longest hypoxic duration of 175 days; it also had the most rapid destratification during fall across years. By comparison, 2018 had the highest average wind speed, with the lowest cumulative radiation, and a somewhat higher O_2_ depletion rate during fall.

**Figure 6.**
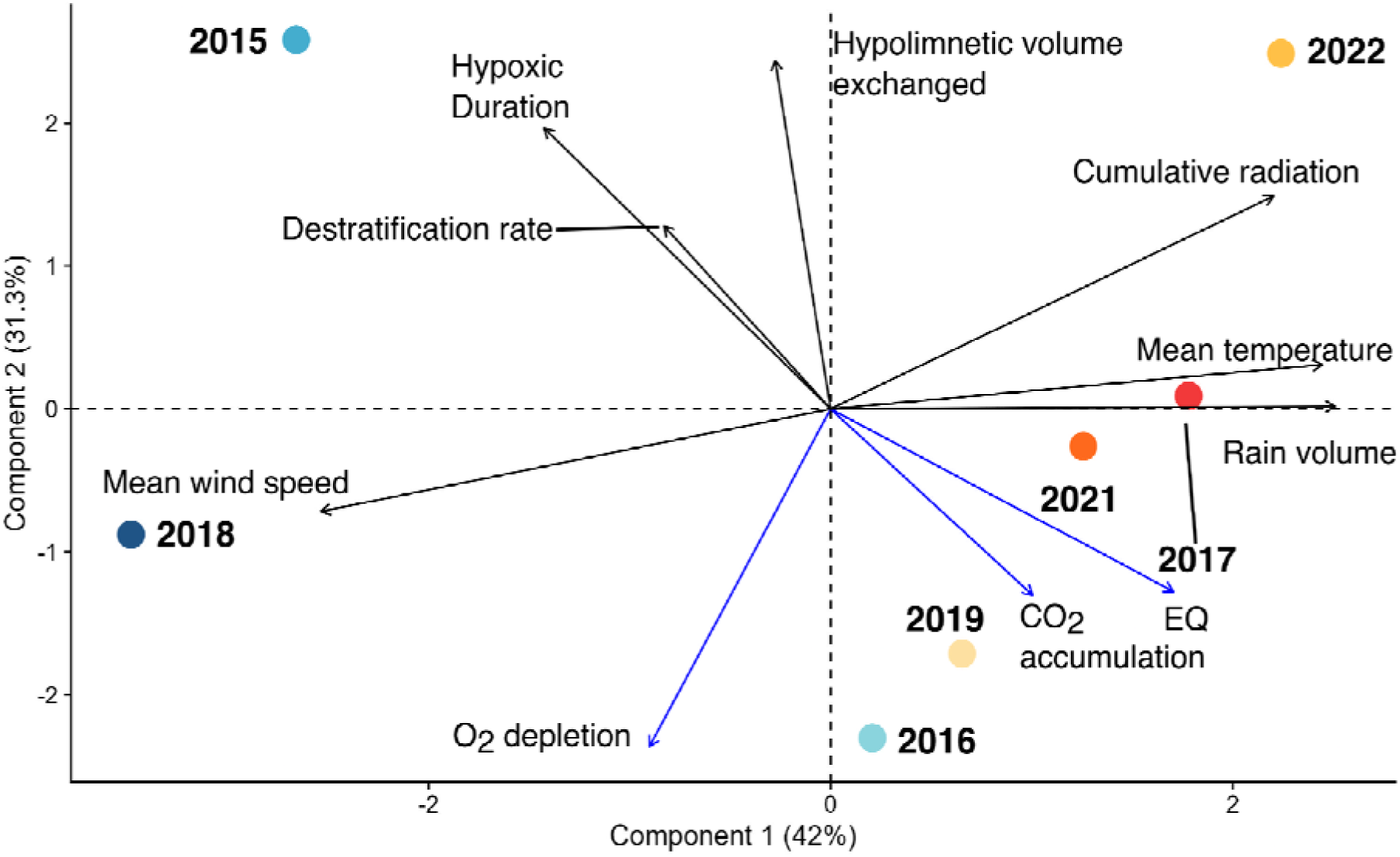
Principal component analysis biplot of years relative to CO_2_ accumulation, O_2_ depletion, EQ, with environmental variables. (mean air temperature during fall, cumulative volume of rain during fall, cumulative radiation during fall, mean wind speed during fall, rate of fall destratification l, summer hypoxic duration, volume of hypolimnion exchanged during fall). Percentage of variance reported on axes.

Higher EQs in 2017, and 2021 were associated with a higher volume of cumulative precipitation and associated watershed inputs, as well as warmer temperatures (Figure 6; Table 5S). In contrast, 2022 had a lower O_2_ depletion that may have been a function of a much higher cumulative radiation encouraging photosynthesis. Internal processes appear to have influenced the EQ of 2019, when the lake had a shorter period of hypoxic duration and exchanged less of its hypolimnetic volume during the study period resulting in high rates of both CO_2_ accumulation and O_2_ depletion. Compared to other years, 2016 had higher O_2_ concentrations in the metalimnion and a small amount of measurable O_2_ in the hypolimnion (Figure 7S), resulting in the highest CO_2_ accumulation rate, likely from continued internal aerobic heterotrophic processing.

### Contrasting CO_2_ lake content

In all years, prior to destratification, the lake contained a certain amount of CO_2_ that was yet to exchange with the surface. We tried to estimate the total amount of CO_2_ that exchanged with the surface as the lake layers broke down and mixed between the cold-dry 2015 and hot-wet 2017 seasons, years that represented two extremes in terms of environmental conditions, and where there was sufficient buoy data to carry out the calculation. This accounting was achieved by estimating the total cumulative amount of CO_2_ in the volume exchanged with the surface volume from adjacent sub water layers during destratification using a mixing approach, and summing the values over time. We found that there was over two times more CO_2_ that exchanged with the surface during the fall of the 2017 hot-wet year (228 kg CO_2_) versus the 2015 cold-dry one (109 kg CO_2_) and that this doubling was a function of a slower, more extended mixing period of 65 days in the former and 32 days in the latter.

## Discussion

This study is, to our knowledge, the first interannual evaluation and characterization of the main drivers influencing surface water O_2_-CO_2_ departures of a temperate freshwater lake during fall. Rather than define the start of the fall season by a calendar date, we used the start of leaf colour change in the forest surrounding the lake. We found that the timing of this phenological trait co-occurred remarkably well with the onset of lake destratification, despite the two being independent indicators of the changing season. This shows that the slowing of evapotranspiration in the forest, which leads to greater external inputs of water and dissolved gases to lake surface waters from the catchment, occurs at the same time as the mixing of hypolimnetic layers into lake surface waters, which contributes internal inputs of dissolved gases. Specifically, we observed strong CO_2_ accumulation and O_2_ depletion in the surface waters during fall each year, compared to those in summer, marking the beginning of a catchment-scale environmental shift across the watershed. In terms of individual metabolic gas dynamics, we observed considerable variability in surface water CO_2_ accumulation rates across years during fall, whereas surface water O_2_ depletion rates were more consistent, with observed rates in six of seven study years being statistically similar to one another. This suggests that variability in CO_2_ accumulation controls the observed difference in paired O_2_-CO_2_ departure signals. The ecosystem quotients (EQs) among years were also largely related to different environmental conditions during the fall, albeit not solely. The variability in departure dynamics during this shoulder season, in contrast to seemingly more stable summer patterns in this lake, likely reflects interannual differences of within lake metabolic processing at depth, as well as the climatic conditions dominating hydrologic processes, making fall a more integrative moment of whole lake ecosystem metabolism.

The higher concentrations of surface water CO_2_ during fall are partially explained by the well-established pattern of hypolimnetic CO_2_ export that occurs during water column mixing in northern dimictic lakes (Kelly and others 2001; Huotari and others 2009; Vachon and del Giorgio 2014). However, our results suggest increased hydrologic inputs as a function of lower catchment water retention capacity, combined with comparatively warmer fall air temperatures, which slowed the rate of destratification that likely extended the contributions of internal processing, best explained years with higher surface CO_2_ accumulation rates. Lakes can receive large amounts of CO_2_ from surrounding catchment soils that are not only strongly supersaturated, but carry substantial amounts of allochthonous organic matter that can further fuel within lake CO_2_ production and corresponding O_2_ respiration via mineralization (Finlay and others 2009; Stets and others 2009; Tranvik and others 2009). Measured CO_2_ concentrations in upland and riparian zone groundwaters in the Lac Croche catchment were found to be almost nine and three-fold higher, respectively, than those measured in the hypolimnion (1849 µmol L^-1^ in the upland and 610 µmol L^-1^ in the riparian zone versus 211 µmol L^-1^ in the hypolimnion), with no measurable O_2_ (see Table 6S) As such, the multiple large precipitation events that occurred towards the end of fall in 2017 and 2019 (Figure 8S), when catchment water retention and lake thermal stability were comparatively very low, appeared to influence CO_2_ inputs. Large storm events have previously been reported to strongly influence lake properties such as thermal stability, turbidity, and increase subsidies (Humborg and others 2010; de Wit and others 2018; Perga and others 2018), with large consequences for lake metabolism, especially during fall (Vachon and del Giorgio 2014). Thus, our results suggest that increased CO_2_ accumulation and EQ in surface waters in hot-wet years, particularly in 2017, are a consequence of higher influx from the surrounding catchment coinciding with increased precipitation and runoff.

Surface CO_2_ accumulation rates and EQs were considerably lower in the colder years (Table 1; Table 2), which also had significantly less precipitation during the fall (Table 5S). In the seven years that were assessed, full lake mixing was only observed in two before the buoy was removed for winter (on October 22, 2015 and October 18, 2018), and were likely driven by higher wind speeds and colder air temperatures for both those years (Butcher and others 2015; Niedrist and others 2018; Woolway and others 2021). Although increases in surface water CO_2_ concentrations by hypolimnetic mixing is well known (Kelly and others 2001; Huotari and others 2009; Vachon and del Giorgio 2014), our results of lower accumulation rates suggest that the supply of CO_2_ from deeper waters was comparatively poor in these cold years. Although the aforementioned lower CO_2_ concentrations in the hypolimnion relative to those of the catchment can partially explain this, it was unexpected given the prolonged period of anoxia in these colder-drier years that would in principle produce more CO_2_ microbially without consuming O_2_ by using other electron acceptors under these prolonged reduced conditions. Bottom waters were nearly anoxic at the onset of buoy deployment for both 2015 and 2018 (Figure 6S and Table 5S), which were also years found to have the larger hypolimnetic volumes (Figure 2). The bottom waters in these cold years were less oxygenated when compared to data reported in other studies working on the same lake (Bogard and others 2017; Massé and others 2019; Pilla and others 2023).

Extended hypoxic duration and prolonged anoxia are of growing concern in temperate boreal lakes, and known to have broader consequences on hypolimnetic chemistry beyond CO_2_ accumulation (Bartosiewicz and others 2019; Jane and others 2023; LaBrie and others 2023; Jansen and others 2024). Our study suggests that changing conditions may also influence lake EQs and potential greenhouse gas emissions at fall turnover in different, and unexpected ways. We anticipated that hypolimnetic waters would bring CO_2_ rich and O_2_ poor waters to the surface during mixing, and although that was the case, we observed that cold-dry years with a higher relative contribution from the hypolimnion experienced lower rates of CO_2_ accumulation suggesting that hypolimnetic CO_2_ accumulation may not have been as important as originally thought in the bottom waters of this oligotrophic, and relatively clear lake.

The difference between the EQs estimated in the fall between more extreme cold-dry years versus the most extreme hot-wet one was nearly double (0.30 in 2015 and 0.38 in 2018 compared to 0.71 in 2017; Table 2), indicating that much less CO_2_ was produced relative to O_2_ consumed in these cold-dry years. EQs from the fall of the cold-wet years are lower than the theoretical respiratory quotient associated with the degradation of allochthonous organic matter (0.7) (Berggren and others 2012). This suggests that some other processes within the lake are either consuming CO_2_ or O_2_ with no concomitant production of the other gas in each respective reaction. One possible explanation is the consumption of CO_2_ during the production of methane. Higher rates of methanogenesis are anticipated in years when hypolimnetic anoxia is prolonged (Bartosiewicz and others 2019), and a recent study has shown that hydrogen-independent CO_2_ reduction methane production may be dominant in a wide range of temperate lakes (Meier and others 2024). Another possible mechanism is O_2_ consumption, exacerbating the depletion rates with no change in CO_2_ via chemical oxidation of reduced species either accumulated in the hypolimnion or diffusing upwards from the sediment. Those species include NH ^+^, Mn(II), Fe(II) and HS^-^ (Steinsberger and others 2019). For instance, in oxic and near neutral pH environments, reduced iron species are quickly oxidized forming Fe(III) (Emmenegger and others 1998; Lau and others 2024) and consume O_2_ in the process. Given the extended hypoxic duration in cold-wet years combined with the capture of complete mixing, it is possible that a combination of these processes resulted in the lower EQs in 2015 and 2018. These aforementioned processes likely occurred in all years but perhaps with lower intensity in years when hypoxic duration was shorter, leading to less accumulation of reduced species and to a signal masked by greater CO_2_ inputs from the catchment. The hypolimnion represented at most 9% of the total lake volume among years, as such it’s likely that only cold-dry years that experienced less cumulative rain during fall (<20% of lake volume, Table 5S), captured a signal dominated by the gas signatures of the hypolimnion.

The intermediate year of 2016 stood out with a much higher fall EQ, largely driven by higher CO_2_ accumulation rate compared to other years, and a somewhat higher O_2_ depletion rate in surface waters that we suspect was likely driven by a difference in internal processing. We suggest that these higher values may be a function of the thicker and more extensive layer of the metalimnetic O_2_ peak observed in 2016 compared to other years (Figure 7S). Although both biological and physical properties are responsible for creating the conditions allowing for a metalimnetic O_2_ maximum (Wilkinson and others 2015), the thick peak observed in 2016 was most likely biological, a phenomenon known to occur in this lake (Ouellet Jobin and Beisner 2014). The high primary production generally associated with such O_2_ maxima (Stefan and others 1995; Wetzel 2001), and the generally higher oxygen concentrations observed in the metalimnion of this lake overall (Figure 6S) may have enabled considerable oxidation of fresh autochthonous organic matter and continued oxic respiration resulting in a larger CO_2_ pool from deeper waters at overturn. O_2_ depletion rates were also likely highest during 2016 as there was simply more O_2_ deeper in the water column, again suggesting that internal processing influenced the fall EQs. In contrast, 2022 had the lowest O_2_ depletion rates during fall, a third lower than other years. We presume that this may be associated with the high light availability that occurred in fall 2022 at 790 MJ m^-2^ that could have potentially fueled more photosynthesis compared to the average of 466 MJ m^-2^ (±100 MJ m^-2^) among other years. These findings highlight how variations in light and within lake O_2_ patterns can influence the overall integrative metabolic response of the lake, emphasizing the close interplay between physical and biological drivers in influencing interannual variations.

## Conclusions

Disentangling the potential drivers of the interannual differences in fall gas dynamics provides insight into the integrative manner in which many factors influence whole-lake metabolism, both during the preceding open water season and winter as well as how conditions vary in preparation for over winter. In this lake, cold-dry fall seasons had lower surface CO_2_ accumulation rates and lower EQs, partially explained by lower watershed inputs combined with a generally longer hypolimnetic hypoxic duration. These lower EQs may have been a function of higher methanogenesis in the hypolimnion (Bartosiewicz and others 2019) expressed as lower CO_2_ accumulation relative to O_2_ loss during overturn. Higher CO_2_ accumulation rates occurred in fall seasons when the lake presumably received more CO_2_ rich waters from its catchment combined with a more gradual input of CO_2_ from bottom waters because of a slowed overturn, which enabled more internal heterotrophic processing at depth through higher aeration and terrestrial subsidies. As such in years where surface concentrations increased mainly from the hypolimnetic inputs, the lake acted as a chimney during fall whereas in those years where it received more rain, the lake acted as a pipe as well as a processor (de Wit and others 2018). Twice as much CO_2_ apparently reached the lake surface when cumulative rainfall was highest and the destratification period was prolonged. Conditions that result in changes to within lake primary production, either as a substantive metalimnetic peak during the summer or potentially a fall bloom also coincided with very different EQs and surface gas dynamics. The interannual variability in gas dynamics and EQs during fall observed here suggests that the metabolic fingerprint of the lake in different years may be largely determined by higher relative amounts of autochthonous C, terrestrial subsidies or inputs of highly processed hypolimnetic organic matter. We suggest that these variable C conditions suggested by the EQs would impact winter food webs (Hébert and others 2021), and under ice metabolism differently that should be further explored. In summary, our study highlights that CO_2_ accumulation and O_2_ depletion rates in lake surface waters as well as metrics of gas departure during fall provide an integrative measure of lake metabolism. As such, a better understanding of the drivers leading to the interannual variability in fall gas dynamics is critical to better predict the impact of future climate change on regional C budgets and potential emissions.

## Acknowledgements

We gratefully acknowledge P. del Giorgo from Université du Québec à Montréal who provided access to the buoy data collected as part of the CarBBAS (Carbon Biogeochemistry in Boreal Aquatic Systems) programme funded by NSERC and Hydro-Québec, which also supported AP who oversaw buoy operations. We would like to thank C. Bélanger, D. Bélanger, L. Galantini, S. Ouimet, S. Shousha who helped either in the lab, field, with model setup, MATLAB scripts or in discussions. We also thank L. Tranvik for feedback. We are grateful to the staff at the Station de Biologie des Laurentides for their logistical support, and access to the PhenoCam information. This work was funded by a CRC Tier 1 in Aquatic Ecosystem Science and Sustainability and NSERC Discovery grant to RM, and a Sentinel North CFREF research chair in Aquatic Geochemistry and NSERC Discovery grant to RMC. BB was partially supported by funding from the Groupe de Recherche Interuniversitaire en Limnologie (FQRNT Strategic cluster). This work was carried out on the traditional lands of the Anishinabiwaki and Omàmiwinini (Algonquin) First Peoples. We are grateful to learn on and learn from these lands.

## Data Availability

Data used for this study are available on the MetaGRIL at https://doi.org/10.5683/SP3/IN5YHU. Scripts for data analysis, figure creation and model are available on GitHub at : https://github.com/blanchette31/Masters

