## Supplemental information for "Interannual surface water CO_2_ and O_2_ dynamics during fall in a small headwater lake"

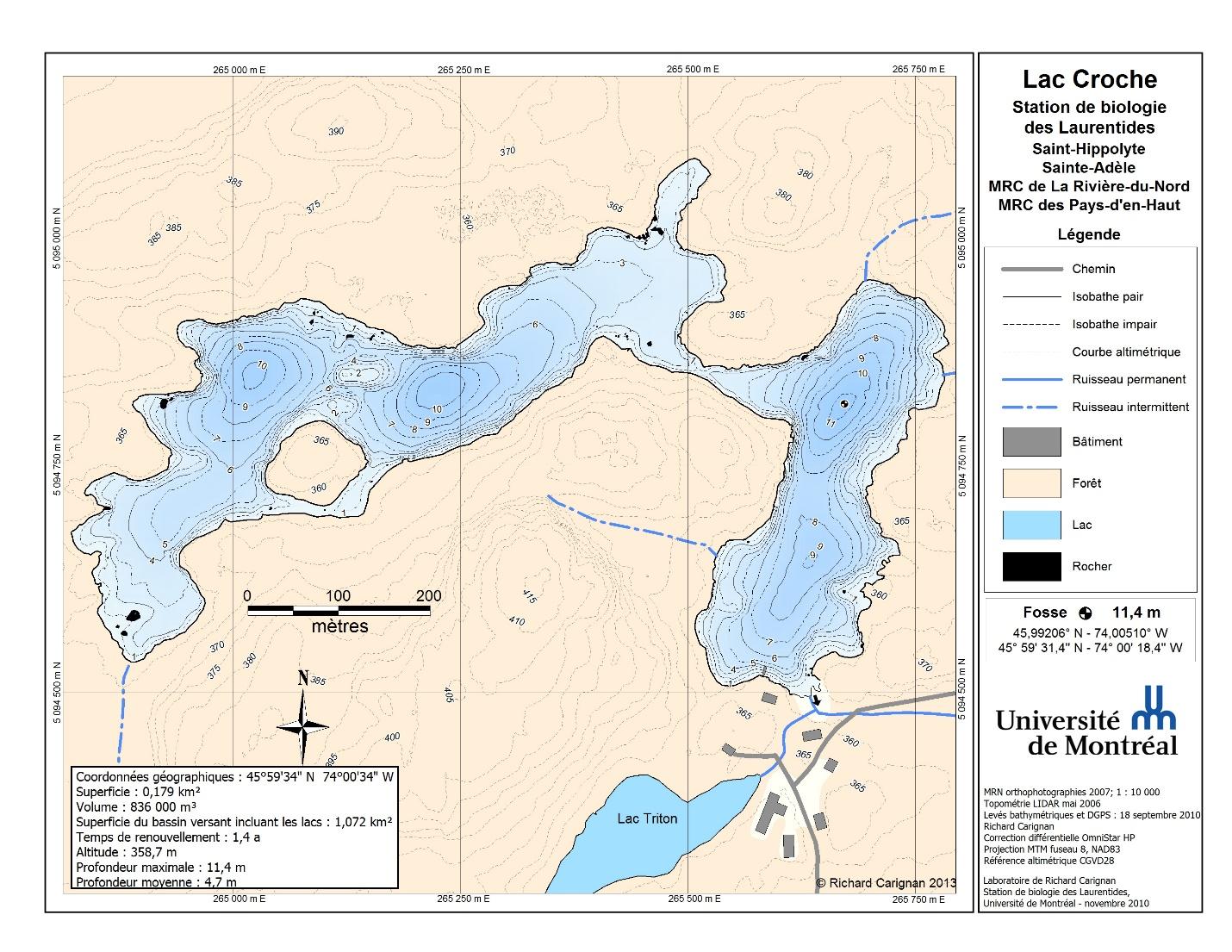

**Figure 1S.** *Bathymetric map of Lac Croche.* The deepest point of the lake (eastern basin) is identified with a white and black circle (From Carignan, 2010). The deepest point of the lake also represents the location of the buoy.

**Table 1S.** *Generalized least square model comparisons for ΔCO_2_ accumulation rates.* Comparison between a model with an autocorrelative structure (auto-regression type 1, AR1) and one without (NO AR1). AIC is used to determine best model fit. The AR1 model was selected for all years as it minimizes AIC.

| Year | Model | Slope | AIC |
| --- | --- | --- | --- |
| 2015 | NO AR1 | 0.40 | 189 |
|  | AR1 | 0.55 | 165 |
| 2016 | NO AR1 | 1.30 | 181 |
|  | AR1 | 1.41 | 176 |
| 2017 | NO AR1 | 0.83 | 363 |
|  | AR1 | 0.81 | 321 |
| 2018 | NO AR1 | 0.40 | 197 |
|  | AR1 | 0.37 | 193 |
| 2019 | NO AR1 | 0.81 | 61 |
|  | AR1 | 0.69 | 62 |
| 2021 | NO AR1 | 0.62 | 167 |
|  | AR1 | 0.59 | 162 |
| 2022 | NO AR1 | 0.61 | 182 |
|  | AR1 | 0.64 | 178 |

**Table 2S.** *Generalized least square model comparisons for ΔO_2_ depletion rates.* Comparison between a model with an autocorrelative structure (auto-regression type 1, AR1) and one without (NO AR1). AIC is used to determine best model fit. AR1 model was selected as it minimizes AIC.

| Year | Model | Slope | AIC |
| --- | --- | --- | --- |
| 2015 | NO AR1 | -1.01 | 326 |
|  | AR1 | -0.99 | 295 |
| 2016 | NO AR1 | -1.16 | 177 |
|  | AR1 | -1.32 | 174 |
| 2017 | NO AR1 | -1.15 | 396 |
|  | AR1 | -1.06 | 335 |
| 2018 | NO AR1 | -1.18 | 227 |
|  | AR1 | -1.16 | 217 |
| 2019 | NO AR1 | -1.27 | 58 |
|  | AR1 | -1.26 | 60 |
| 2021 | NO AR1 | -1.14 | 180 |
|  | AR1 | -1.01 | 160 |
| 2022 | NO AR1 | -0.64 | 211 |
|  | AR1 | -0.62 | 202 |

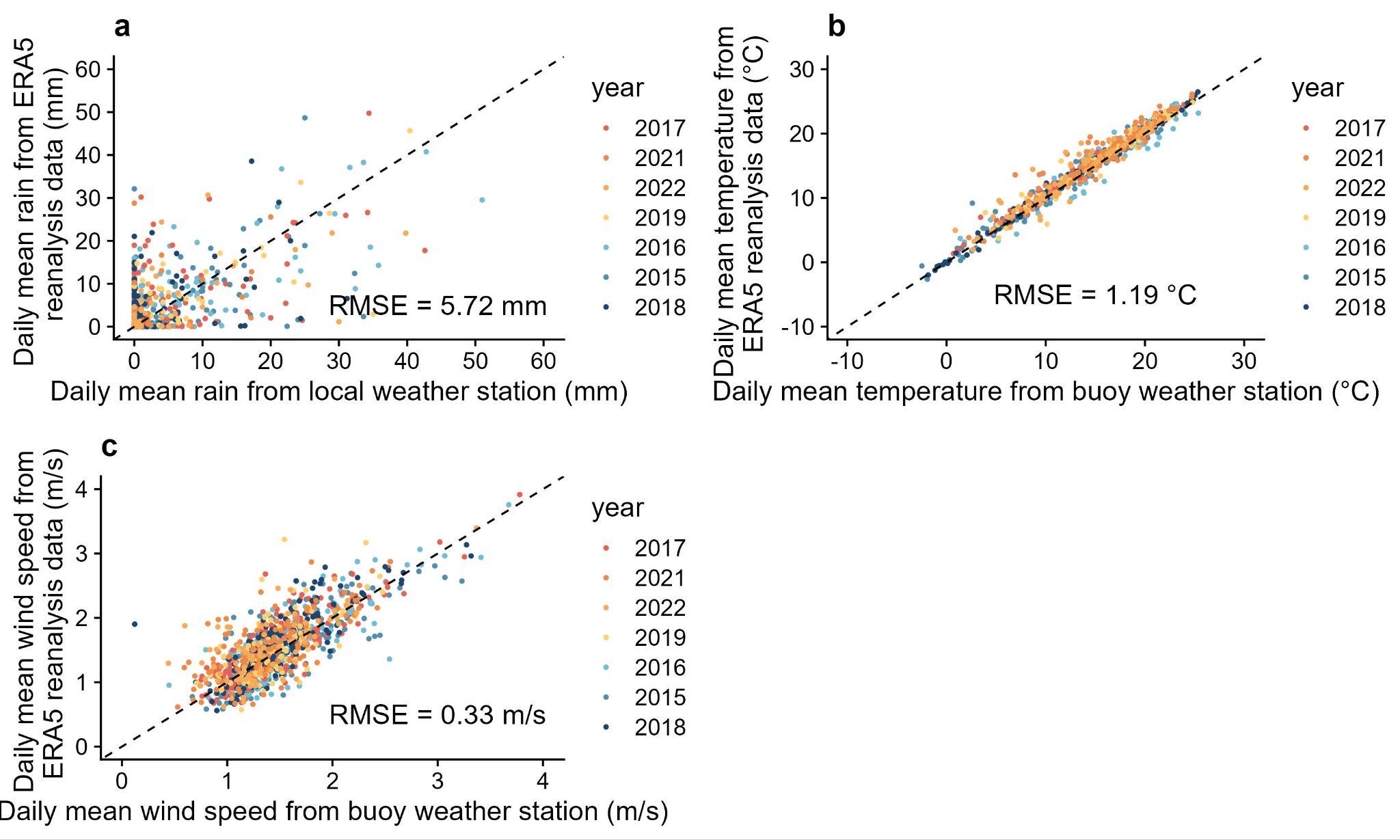

**Figure 2S.** *Validation between local and ERA5 weather data.* Figure a) compares the daily precipitation between the local data and ERA5 modelled data. Figure b) and c) compare the daily average air temperature and wind speed (m s^-1^) respectively. A correction factor of 0.53 was applied to ERA5 data to improve fit with data from the weather station on the buoy.

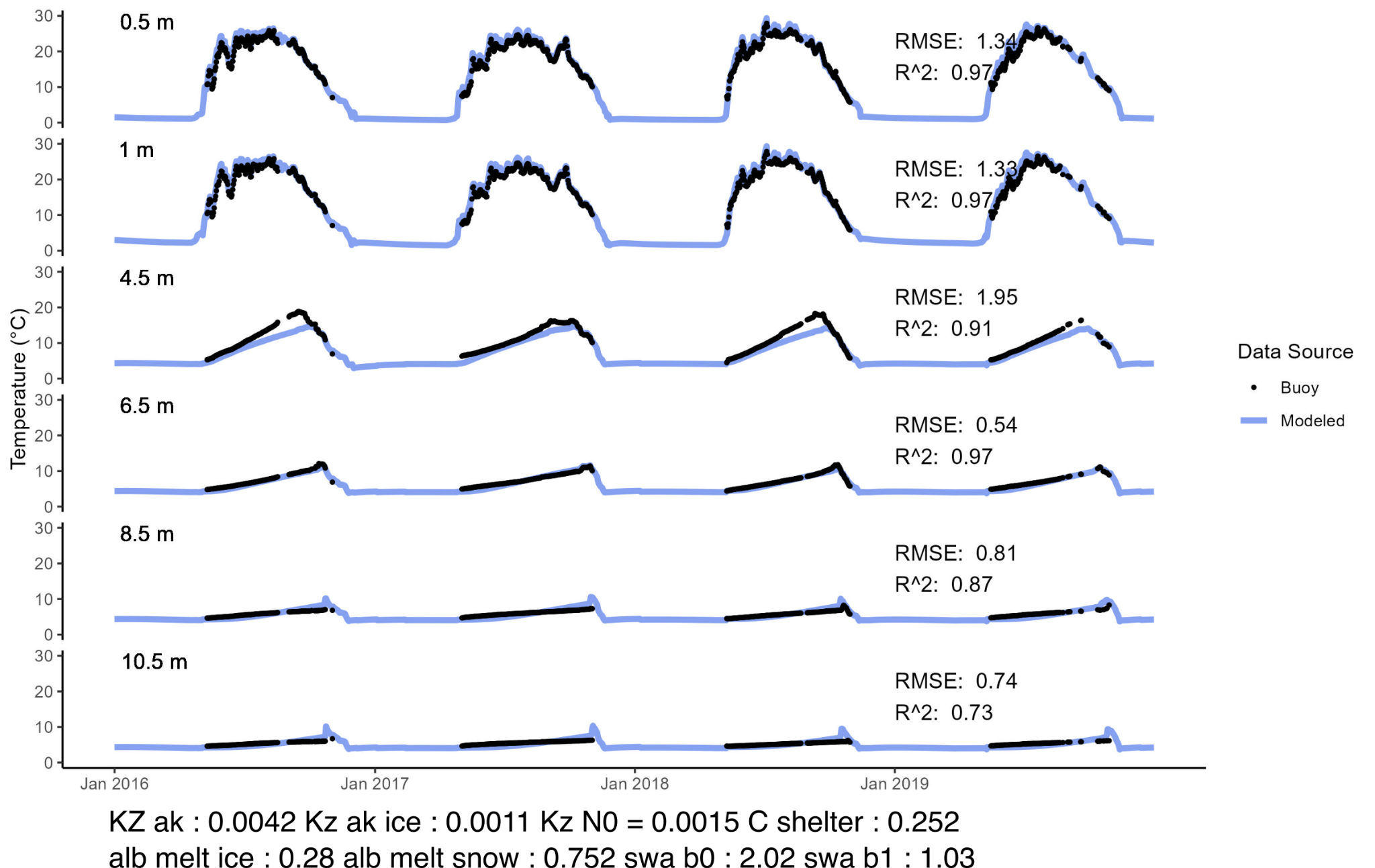

**Figure 3S.** *Model water temperature calibration run 2016-2019 at specified depths (0.5m, 1.0m, 4.5m, 6.5m, 8.5m, 10.5m).* Blue line represents the daily time step temperature prediction while the black dots represent the measured daily average temperature from the buoy. Parameter values for calibration listed below figure.

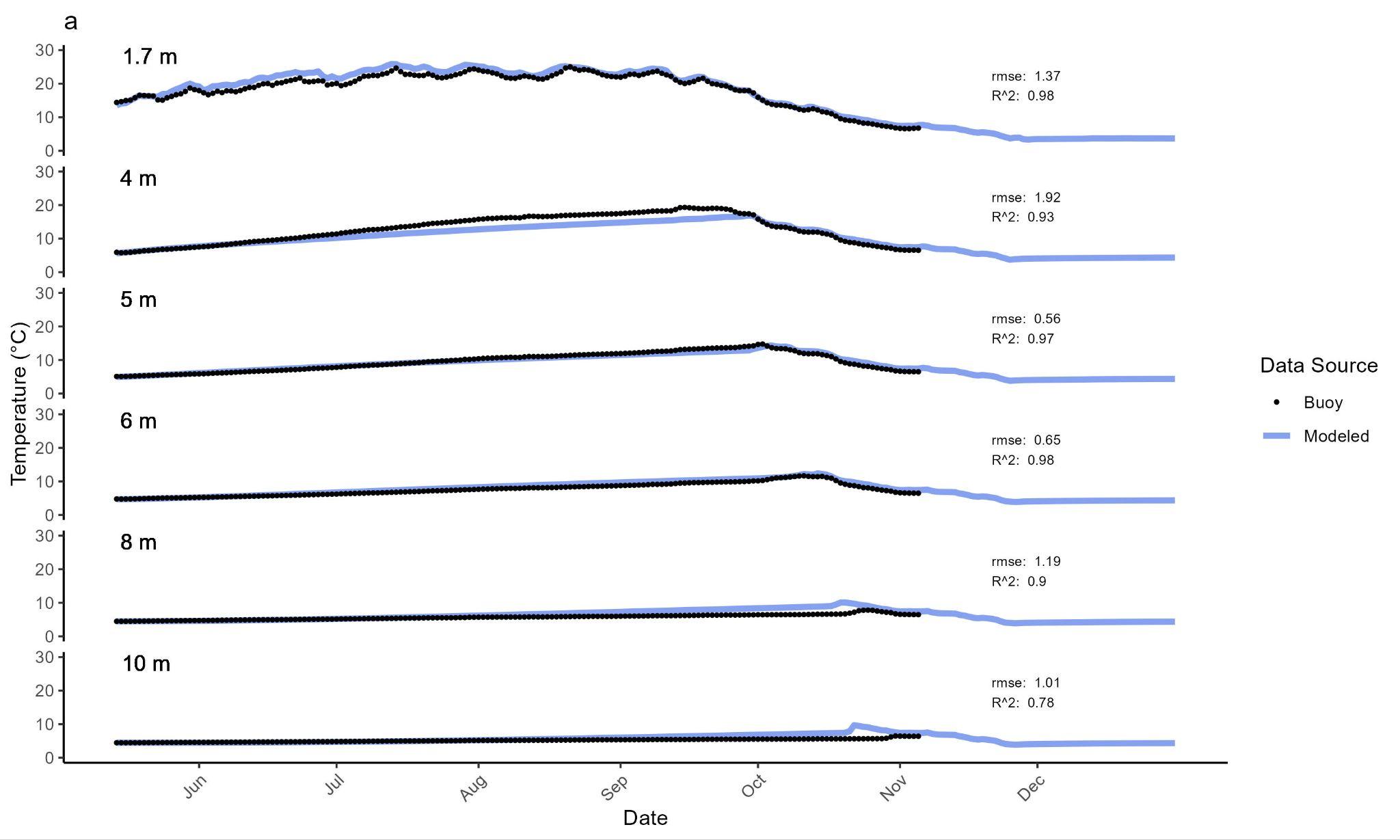

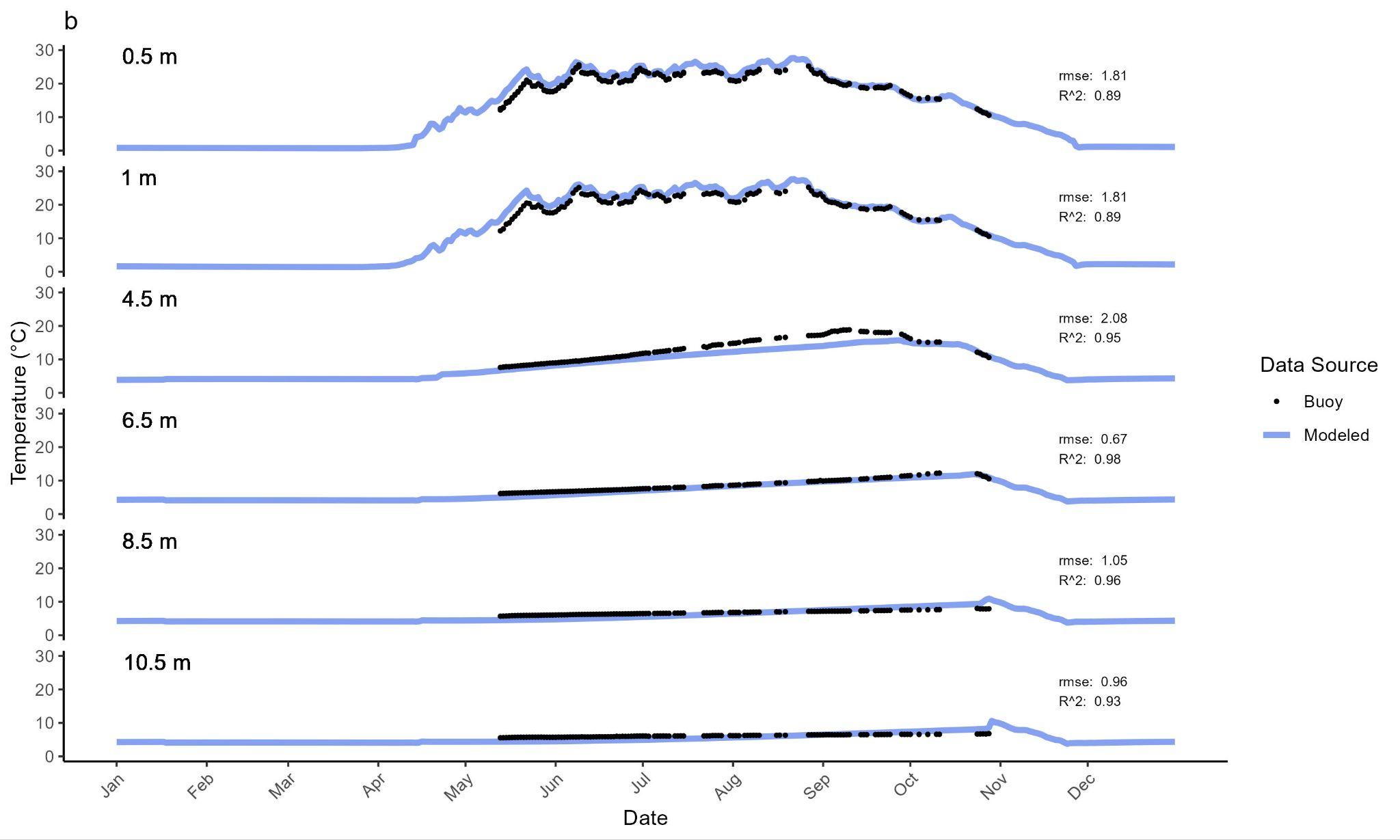

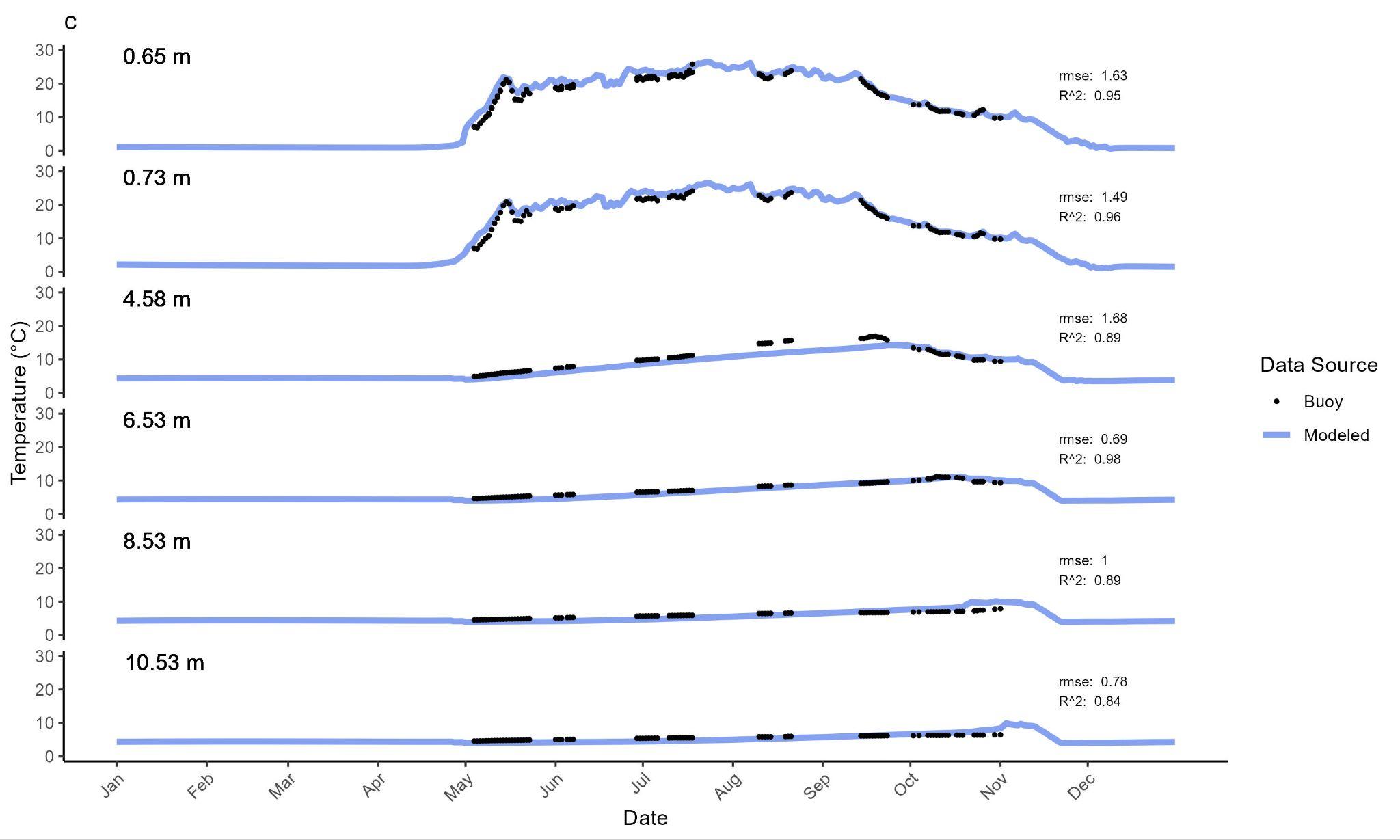
**Figure 4S.** *Model temperature validation for 2015 (a) and 2021 (b) and 2022 (c).* Blue line represents the daily time step temperature prediction while the black dots represent the measured daily average temperature from the buoy.

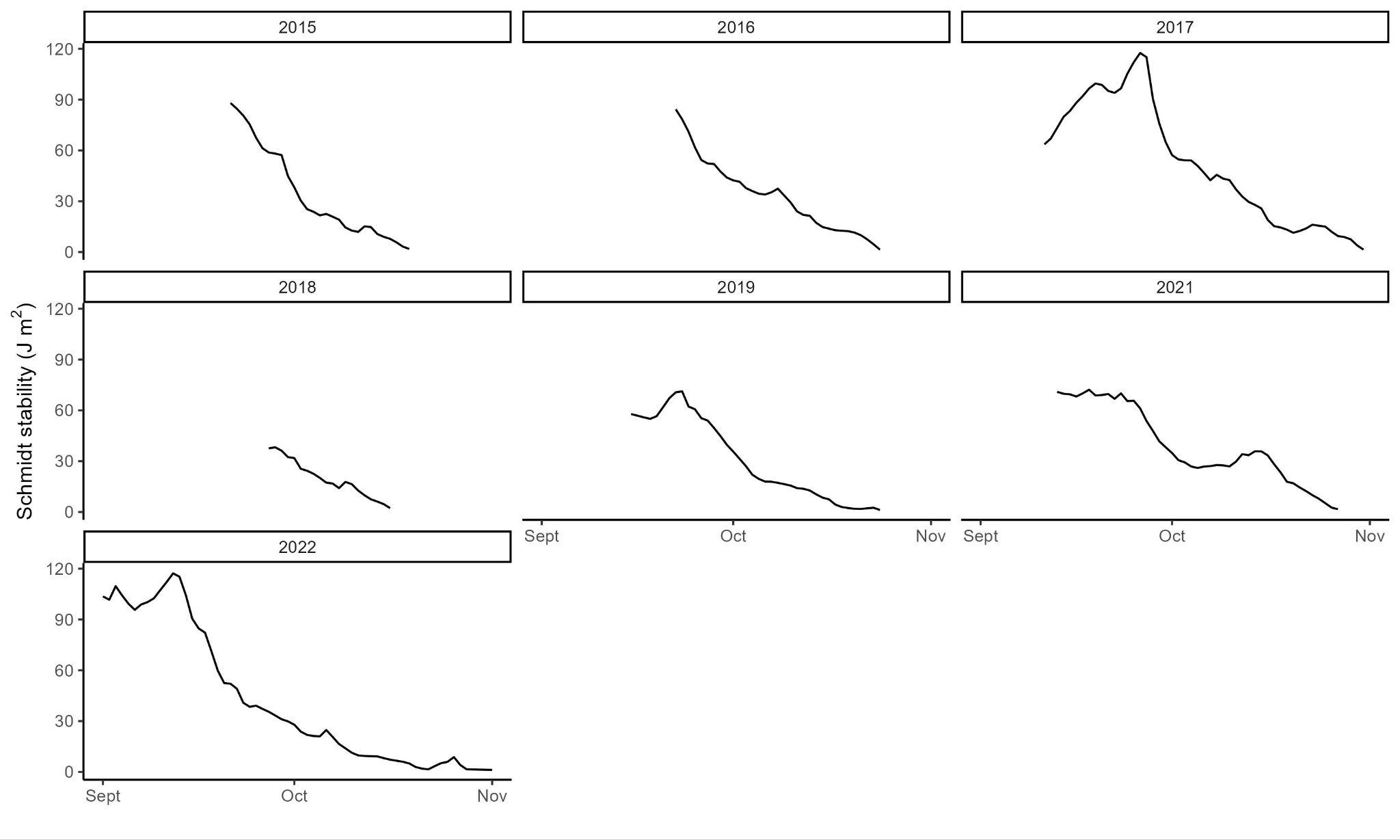

**Figure 5S.** Schmidt stability as a function of time (day of year) from the start of fall to the end of the study period for each year.

**Table 3S.** *Results of pairwise Tukey contrasts of ΔCO2 accumulation slopes.* Only significantly different pairs (p < 0.05) are reported here.

| Pair | Estimate | Standard error | t-value | p-value |
| --- | --- | --- | --- | --- |
| 2015-2016 | -235.742 | 38.410 | -6.137 | < 0.001 |
| 2015-2017 | -101.163 | 30.516 | -3.315 | 0.01754 |
| 2016-2017 | 134.580 | 30.879 | 4.358 | < 0.001 |
| 2016-2018 | 260.406 | 39.491 | 6.594 | < 0.001 |
| 2016-2019 | 152.510 | 48.185 | 3.165 | 0.02749 |
| 2016-2021 | 194.531 | 35.772 | 5.438 | < 0.001 |
| 2016-2022 | 192.240 | 33.997 | 5.655 | < 0.001 |
| 2017-2018 | 125.827 | 31.866 | 3.949 | 0.00196 |

**Table 4S.** *Results of pairwise Tukey contrasts of ΔO_2_ depletion slopes.* Only significantly different pairs (*p* < 0.05) are reported here.

| Pair | Estimate | Standard error | t-value | p-value |
| --- | --- | --- | --- | --- |
| 2022-2016 | -136.513 | 47.840 | -2.854 | 0.06528 |
| 2022-2017 | -133.708 | 34.802 | -3.842 | 0.00276 |
| 2022-2021 | -129.805 | 43.092 | -3.012 | 0.04202 |

| Year | Cumulative global radiation summer | Cumulative global radiation fall | Mean temperature summer | Mean temperature fall | Volume rain during fall | Mixing volume | Hypoxic duration | Fall stratification breakdown rate | Mean wind speed fall | Leaf colour change |
| --- | --- | --- | --- | --- | --- | --- | --- | --- | --- | --- |
|  | MJ m^-2^ | MJ m^-2^ | (℃) | (℃) | (% lake volume) | (m^-3^) | (days) | (J m^-2^ d^-1^) | (m s^-1^) | (date) |
| 2015 | 1409 | 459 | 17.3 | 6.89 | 19 | 153936 | 175 | 1.86 | 3.201 | Sept. 21 |
| 2016 | 1403 | 414 | 17.2 | 7.94 | 22 | 111492 | 126 | 1.46 | 3.086 | Sept. 21 |
| 2017 | 1230 | 625 | 15.4 | 12.9 | 31 | 111492 | 130 | 1.56 | 3 | Sept 11 |
| 2018 | 1525 | 296 | 17.6 | 5.17 | 12 | 111492 | 135 | 1.21 | 3.394 | Sept 27 |
| 2019 | 1366 | 489 | 15.6 | 9.16 | 26 | 111492 | 102 | 1.08 | 3.047 | Sept 15 |
| 2021 | 1360 | 510 | 17.2 | 11.5 | 27 | 108721 | 127 | 1.11 | 2.89 | Sept. 13 |
| 2022 | 1204 | 790 | 17.2 | 10.5 | 25 | 151156 | 134 | 1.26 | 2.789 | Sept. 1 |

**Table 5S.** *Climatic and physical variables as potential drivers for surface water metabolic gas dynamics.* Fall is defined as the leaf loss period of each year.

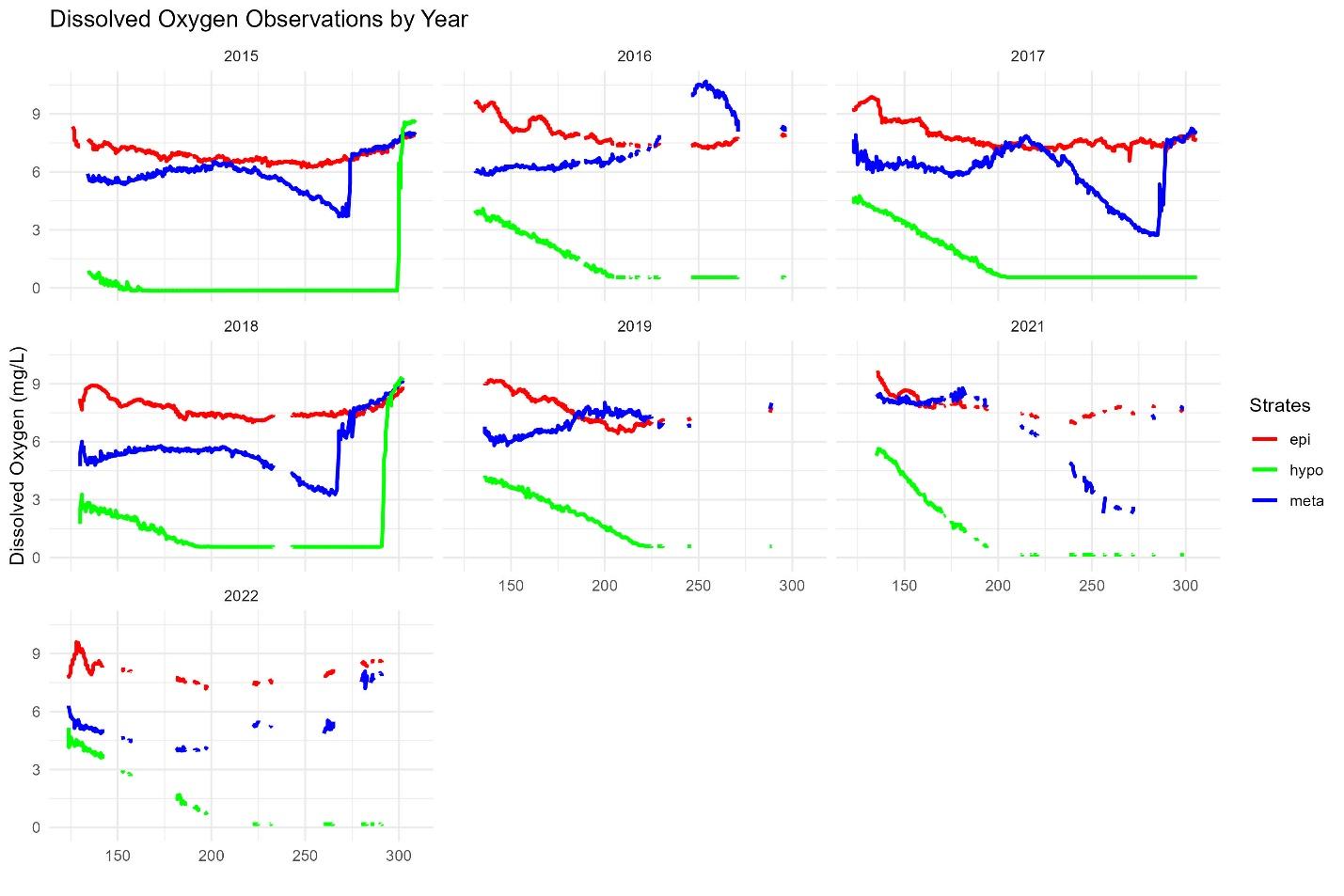

**Figure 6S.** *O_2_ concentrations in mg L^-1^ at different depths over time (day of year) across years.* Red lines are epilimnetic O_2_ concentrations, blues lines are metalimnetic and green lines are hypolimnetic O_2_ over time. Depth of sondes for each strata is 0.5m for the epilimnion, 5.5m for the metalimnion and 8.5m for the hypolimnion.

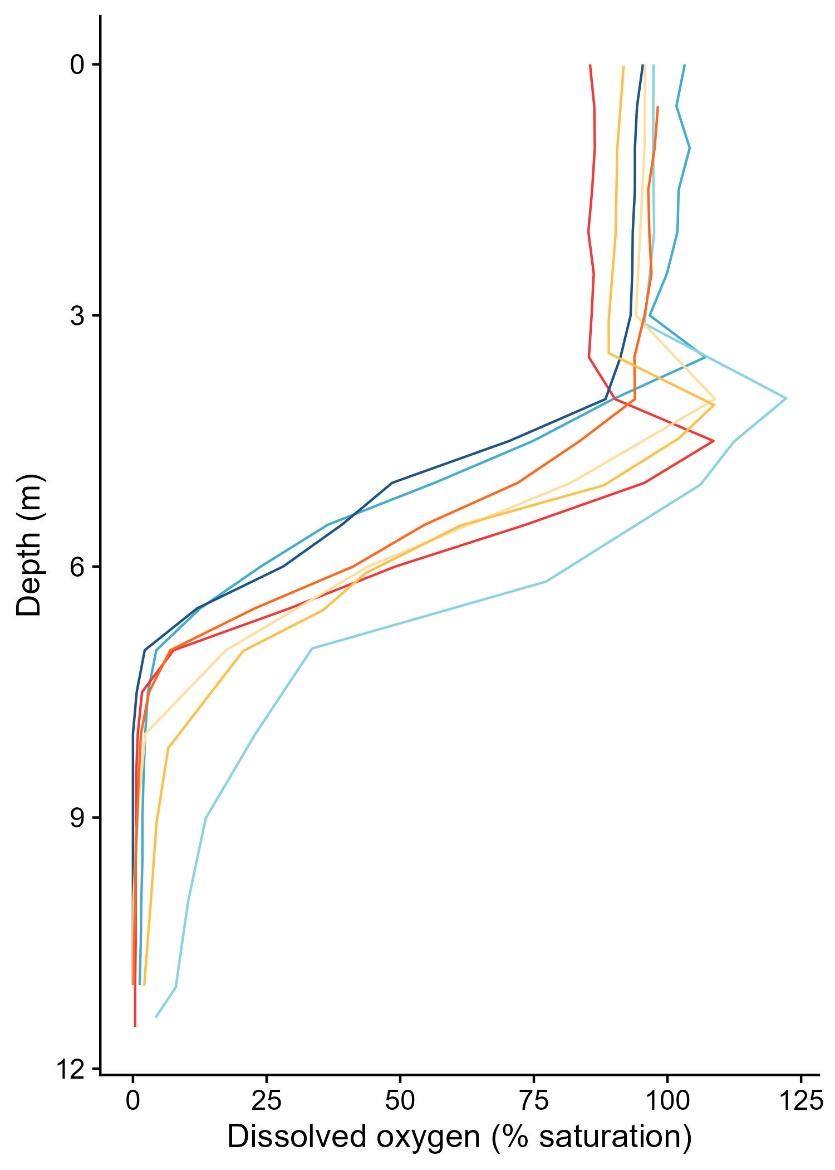

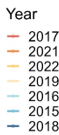

**Figure 7S.** Lac Croche dissolved oxygen profiles during the late summer (~August 18^th^) of each year.

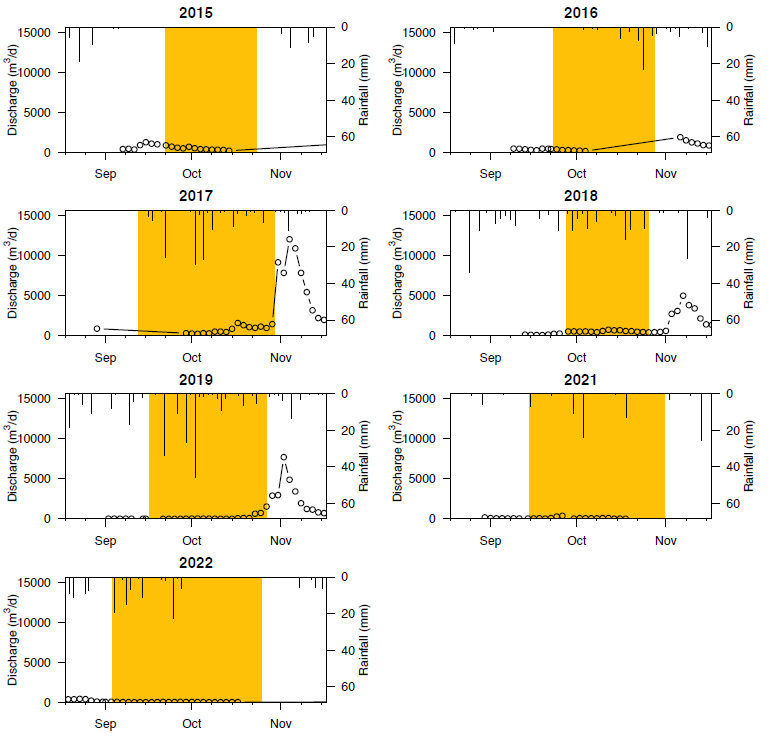

**Figure 8S.** *Daily precipitation and discharge over time across years.* Yellow box represents fall for each year. Period shown is from August 18^th^ to November 14^th^ of each year.

**Table 6S.** Mean CO_2_ concentrations in µmol L^-1^ of samples taken on November 11^th^, 2023 in the forested, riparian zone and stream sites and on November 11^th^ 2022 for the hypolimnion.

| Site | CO_2_ concentration (µmol L^-1^) |
| --- | --- |
| Forested | 1849 |
| Riparian zone | 610 |
| Stream | 357 |
| Hypolimnion | 211 |
